# *Plasmodium falciparum* immunodominant IgG epitopes in subclinical malaria

**DOI:** 10.1101/792499

**Authors:** Isabel G. Azcárate, Patricia Marín-García, Paloma Abad, Susana Pérez-Benavente, Estela Paz-Artal, Pedro A. Reche, Julius N. Fobil, José M. Rubio, Amalia Diez, Antonio Puyet, José M. Bautista

## Abstract

Incomplete non-sterile immunity to malaria is attained in endemic regions after recurrent infections by a large percentage of the adult population, who carry the malaria parasite asymptomatically. Although blood-stage *Plasmodium falciparum* rapidly elicits IgG responses, the target antigens of partially protective and non-protective IgG antibodies as well as the basis for the acquisition of these antibodies remain largely unknown. We performed IgG-immunomics to screen for *P. falciparum* antigens and to identify epitopes associated with exposure and clinical disease. Sera from malaria cases identified five prevalent antigens recognized by all analyzed patients’ IgGs. For further epitope mapping, peptide microarrays designed to cover their sequences were probed with a set of 38 sera samples from adult individuals of an endemic malaria region in Ghana. Eight 20-mer peptides with the highest affinity and frequency of recognition among the population were subsequently validated with 16 sera from the same region, segregated into patients with positive or negative subclinical detection of *P. falciparum*. Significant binding specificity for two immunodominant antigenic regions was uncovered within the START-related lipid transfer protein and the protein disulfide isomerase PDI8. These 20-mer peptides challenged with sera samples from children under 5 years old displayed specific IgG binding in those with detectable parasitemia, even at subclinical level. These results suggest that the humoral response against START and PDI8 antigens may be triggered even at submicroscopic parasitemia levels in children and may eventually be used to differentially diagnose subclinical malaria in children.

**Significance:** Malaria in Africa is a leading cause of morbidity and mortality. The reservoirs of the malaria parasite are asymptomatic patients who carry it subclinically. Identifying the parasite antigens and its fragments that trigger the most common immunity response by immunoglobulin G that partially protect people can have profound implications for both, development of a malaria vaccine and diagnosis of the subclinical parasite carriers. Antigen discovery and mapping, validated with sera from subclinical carriers, showed that immunoglobulin G responses in children against parasite’s START and PDI8 may eventually be used to differentially diagnose non-infected from subclinical cases. Furthermore, anti-START and anti-PDI8 endemic immunodominance provides association of these antigens with long-term acquired immunity and immune evasion to malaria.

## Introduction

Endemic *Plasmodium falciparum* malaria causes million clinical cases and hundreds of thousand deaths worldwide (1), although the global disease toll probably exceed these numbers (2, 3). Most deaths occur in sub-Saharan Africa (90%) and in children under 5 years old (70%) (1). In non-endemic areas, malaria could also be a public health threat due to international tourism and migration to and from endemic areas, which has led to the occasional re-emergence of this parasitic disease as imported infection (4). In-housing vector control, elementary clinical diagnosis and therapeutic drugs are the only available management strategy for both malaria prevention and treatment in endemic regions. As a necessary step towards malaria elimination neither vaccine nor diagnostics of subclinical reservoirs are widely available. After repeated exposure to the parasite, those who survive malaria early in life eventually acquire resistance to the disease (5), but the mechanisms that underlie this immunity remain poorly understood (6). Landmark studies in the 1960s showed that purified IgG from malaria-immune adults transferred to acute-malaria infected children leads to reduction of both fever and parasitemia (7), thus indicating that antibodies against *P. falciparum* proteins play a critical role in controlling the blood stage of the infection. However, it is still unknown which of the 5,300 proteins presently annotated in the *P. falciparum* genome (PlasmoDB, www.plasmodb.org) elicit the production of protective antibodies (8). In addition to allelic variations of MHC molecules, the degree of antigen exposure, antigen abundance and host immunodominance impede that all possible antigens could be recognized by natural immune responses (9). With regard to host immune detection of the *P. falciparum* parasite, pioneer studies showed that a large number of antigens are recognizable and dispersed amongst a large fraction of the proteome (10, 11). Among these, specific epitopes that do induce a response upon immunization with whole native antigens are defined as dominant (12). Although most of the putatively exposed parts of the Ag surface could be subject of recognition by antibodies, the selection mechanism by which only certain antigen regions become B-cell epitopes is not fully understood (13). The accurate identification of B-cell epitopes constitutes a basis for development of antibody therapeutics (14), peptide-based vaccines (14-16), and immunodiagnostic tool (17).

Previous studies in areas endemic for malaria have been carried out to identify a correlation between malaria immunity and *P. falciparum*-specific antibodies. These studies determined antibody reactivity against either a few selected proteins available after cloning (18) or a wider selection of the *P. falciparum* proteome (up to 23%) by means of microarray technology (11). Scanty information is available about the actual immunodominant B-cell epitopes from the identified antigens. Based on the spatial structure, epitopes can be categorized as a continuous (linear or sequential) and discontinuous (nonlinear or conformational); in the latter case amino acid residues are in close contact due to the three-dimensional conformation (16, 19). The amount of amino acid sequence in native proteins required for the correct folding of a discontinuous epitope of B cells is always greater than 20 amino acids and can reach up to 400 amino acids. Using a less stringent definition for continuity, it has been proposed that the majority of discontinuous epitopes (over 70%) are composed of 1–5 linear segments of lengths of 1–6 amino acids (20). Currently, the Immune Epitope Database (http://www.iedb.org/home_v3.php) contains above 700 epitopes from 63 antigens of *P. falciparum*. Most of them have been identified in commonly identified antigens, i.e. MSP proteins (21-23), circumsporozoite protein (CSP) (24, 25) or ring-infected erythrocyte surface protein (RESA) (26) among others. Notwithstanding all these data, correlation between IgG response for the outlined *P. falciparum* proteins and malaria immunity have not been firmly established, suggesting that antibodies against these proteins might not play a role in protective immunity or, more likely, that IgG against single-parasite proteins are insufficient to confer protection. In addition, the identification of B-cell epitopes can also be used in the development of diagnostic tests by immunodetection approaches (27). These data highlight the importance of increasing the repertoire of antigens and epitopes naturally expressed in Plasmodium by different strategies.

In an effort to expand the search of *P. falciparum* antigens and immunodominant epitopes, we first analyzed immunoproteomic profiles shown in sera from patients with imported malaria. These patients living in a non-endemic region were not periodically re-infected after other continuous *P. falciparum* infections, but occasionally when they visit transiently their home country of endemic malaria. Thus, their humoral immune status may hypothetically reflect a selection for pre-immunogenic anti-*P. falciparum* IgG antibodies, eventually directed against epitopes that suppressed acute symptoms. The identification of IgG-reactive antigens in sera from imported malaria patients with a single recent restricted infection but with different levels of parasitemia would therefore be particularly informative, as this could allow for the identification of immunoreactive antigens that favour subclinical malaria and therefore may be relevant for both immunisation and diagnosis. In this work, the humoral response in imported malaria patients was analyzed for the identification of potential highly immunoreactive antigens and mapping of immunodominant epitopes using peptide microarrays. Subsequently, the specific IgG response to a selection of these epitopes in adult and child sera from a malaria endemic Ghanaian population was quantified, validating the antigens found and suggesting a possible correlation with the stage of development of the infection.

## Results

### Immunoproteomics of *P. falciparum* blood stage antigens

The sera of 19 imported malaria patients showed *P. falciparum* infection through parasitological and serological analyses. Five of these patients showed clinical symptoms and were classified as imported clinical malaria (ICM) with high parasitemia. The remaining 14 patients were identified during routine medical checks and were classified as imported subclinical malaria (ISC), since no clinical symptoms of malaria were observed in addition to a submicroscopic parasitemia. Four additional sera from individuals who had never been in malaria-endemic regions were used as control. Mean values of specific anti-*P. falciparum* IgG antibodies quantified by indirect ELISA did not show significant differences between both patient groups, although very large intragroup variability was observed in the ISC group (Figure S1A). No specific antibodies against *P. falciparum* were detected in unexposed controls (not shown). Although the total amount of anti-*P. falciparum* antibodies remained fairly constant regardless of clinical symptoms, differences in antibody avidity were observed between serum samples of the two stages of infection; namely ISC and ICM (Figure S1B), suggesting that a different collection of anti-*P. falciparum* IgG species may exist in sera of patients with very low parasitemia (higher avidity) compared to patients with clinical symptoms (lower avidity). To further identify differential IgG reactive antigens, detailed immunoproteomics was performed. Thus, the total *P. falciparum* intraerythrocytic proteome obtained in 2D-PAGE from a clinical isolate, originally causing subclinical malaria (28), showed 862±14 spots within the range at 3-11 isoelectric point and 3-260 kDa molecular weight (Figure S2). The corresponding 2D immunoblots using sera from imported malaria cases revealed different immunoreactivity patterns for each group of patients (Figure S2). Immunoblots obtained with ICM group sera showed a higher number of immunoreactive spots than ISC group sera, suggesting that parasitemia correlates with obtaining a diverse and strong humoral immune response. In addition, ISC sera showed a predominantly acidic protein pattern, in contrast to immunoblots obtained from ICM sera that showed a wide variety of both acid and basic antigens.

The quantification of signals from all samples allowed the selection of 83 immunoreactive protein spots that were present in at least 30% of the samples from each patient group, ICM or ISC. These spots were excised from the reference 2D gel and subjected to MALDI-TOF/TOF analysis and subsequent search in *P. falciparum* databases for protein identification. Although nearly 60% of the removed spots could not be identified (<90 score), the remaining 34 spots (Figure S2) were successfully assigned to 17 proteins, as shown in Table 1, and classified into the main functional classes of the *P. falciparum* proteome (29), resulting in seven of protein fate, five of metabolism, four of cell surface or organelles and only one of transport. Although some differences were observed for antigen reactivity, none of the identified proteins appeared to be significantly associated with clinical or subclinical stages. However, with the expectation of characterizing potential differences in epitope recognition, five antigens were selected based on relative immunodominance (four of which showed a positive identification of >50% by patient sera) and computational analysis of antigen function and recognition by T and B cells (30, 31): Putative protein kinase (Q8I2Q0), Heat shock protein 70 (HSP70-2), Heat shock protein 60 (HSP60) and Star-related lipid transfer protein (START), and the Protein disulfide isomerase 8 (PDI8). Among the five selected proteins, only HSP60 and PDI8 had been reported as antigenic in previous studies (11, 32).

**Table 1.** Immunoproteomics-based identification of *P. falciparum* proteins that showed IgG reactivity with serum samples from malaria patients. Information is included on its function and on the developmental stages where its gene expression and protein have been detected from previous studies (29, 45).

| Spot n. | Protein | Gene name | Accession | Score | ISC (%) | ICM (%) | Total | Localization/Function | Functional category | Gene Expression (All Stages)** | Protein detection in erythrocytic stages *** |
| --- | --- | --- | --- | --- | --- | --- | --- | --- | --- | --- | --- |
| 1,2,3,4 | Putative protein kinase |  | Q8I2Q0* | 485 | 86 | 83 | 85 | Unknown/putative protein kinase | Protein fate | Sexual & Asexual TR (very low) | MR, TR, GM |
| 13,14,15,16 | Heat shock protein 70 | HSP70-2 | Q8I2X4* | 1220 | 71 | 100 | 80 | Cytoplasm/Stress response protein binding | Protein fate | Sexual & Asexual TR | ALL |
| 17,18,19 | Heat shock protein hsp70 homologue | HSP70-3 | Q9GUX1 | 1110 | 71 | 67 | 70 | Cytoplasm/Stress response protein binding | Protein fate | Sexual (mostly) Early-Mid TR | ALL (TR predominant) |
| 7,8,9,10,11,12 | Heat shock protein 60 | HSP60 | Q8IJN9* | 1310 | 64 | 83 | 70 | Cytoplasm/Stress response protein binding | Protein fate | Sexual & Asexual Late RG-Mid TR (low) | ALL (TR predominant) |
| 20,21 | Heat shock protein 70 | HSP70 | Q8IB24 | 819 | 57 | 50 | 55 | Cytoplasm/Stress response protein binding | Protein fate | Sexual & Asexual RG-TR | ALL (TR predominant) |
| 5,6 | Star-related lipid transfer protein | START | Q8I298* | 509 | 50 | 67 | 55 | Integral membrane/Phospholipid transfer | Transport | Asexual ALL (low) | MR, GM |
| 27 | Plasmeprin I | PMI | Q7KQM4 | 881 | 50 | 50 | 50 | Vacuole/Aspartic endopeptidase | Metabolism | Asexual (low) | MR, GM, TR (TR predominant) |
| 28 | Merozoite surface protein 9 | MSP9 | C5HEE4 | 435 | 43 | 67 | 50 | Membrane/erythrocyte invasion | Surface or organelles | Asexual (predominant) Late TR-Early SC (high) | MR, TR |
| 29 | Merozoite surface protein 1 | MSP1 | A5A7C2 | 889 | 50 | 33 | 45 | Membrane/erythrocyte invasion | Surface or organelles | Asexual Mid TR-Late SC (very high) | ALL (MR predominant) |
| 33 | Dihydrolipoamide acetyltransferase component of pyruvate dehydrogenase complex | BCKDH-E2 | O97227 | 707 | 43 | 50 | 45 | Mitochondrion/acetyltransferase activity | Metabolism | Sexual & Asexual<br>Early TR-Early SC | MR, GM, TR<br>(GM predominant) |
| 30 | Ferredoxin reductase-like protein |  | Q8IBP8 | 279 | 36 | 67 | 45 | Oxidoreductase | Metabolism | Sexual (low) & Asexual<br>TR (low) | SC, GM<br>(Predominant GM) |
| 23,24,25 | Protein disulfide isomerase | PDI8 | C0H4Y6* | 1300 | 43 | 33 | 40 | Endoplasmic reticulum/protein folding | Protein fate | Sexual & Asexual<br>TR | ALL |
| 31 | Dihydrolipoyl dehydrogenase | LPD1 | Q8I5A0 | 1040 | 43 | 33 | 40 | Cytoplasm/Cell redox homeostasis | Metabolism | Sexual & Asexual<br>TR | ALL<br>(TR & GM predominant) |
| 32 | V-Type proton ATPase subunit B |  | Q6ZMA8 | 1090 | 43 | 17 | 35 | Vacuole/ATP hydrolysis coupled proton transport | Transport | Sexual & Asexual<br>Early TR-Early SC | ALL<br>(MR, TR & GM predominant) |
| 26 | Plasmeprin II | PMII | P46925 | 623 | 36 | 33 | 35 | Vacuole/Aspartic endopeptidase | Metabolism | Sexual & Asexual<br>TR (low) | MR, TR, GM<br>(TR predominant) |
| 22 | Heat shock protein 90 | HSP90 | Q8IC05 | 637 | 36 | 33 | 35 | Cytoplasm/Stress response protein binding | Protein fate | Sexual & Asexual<br>RG, TR | ALL<br>(TR predominant) |
| 34 | Rhoptry-associated protein 2 | RAP2 | Q8I484 | 93 | 29 | 33 | 30 | Rhoptry/cell invasion | Surface or organelles | Asexual (only)<br>Early-mid SC | MR, TR, GM<br>(MR predominant) |

\* Proteins selected for epitope mapping
\*\* Data from Malaria Cell Atlas (45). Stage expression is indicated as follows: Ring (RG), Trophozoite (TR), Schizont (SC), All Asexual Stages (ALL)
\*\*\* Data from (29). Stage expression is indicated as follows: Ring (RG), Trophozoite (TR), Schizont (SC), Merozoite (MR), Gametocyte (GM), All Erythrocytic Stages (ALL)

### Mapping linear IgG epitopes with sera from an endemic malaria region

If any of the antigens detected in imported malaria cases are relevant to the maintenance of acquired malaria immunity, sera from asymptomatic individuals in endemic areas can be expected to contain specific antibodies against epitopes in these proteins. To analyze this possibility, random sampling of sera from patients visiting Breman-Asikuma Hospital (Ghana) for reasons other than malaria symptomatology was undertaken. Blood samples were quantified for *P. falciparum* parasitemia and IgG content, and subsequently HLA genotyped. The sera were classified into two groups: adults with subclinical malaria (ASC) with low parasitemia level (<0.1%); and adults with no parasitemia (ANP), and thus no having malaria. Both groups were further subdivided by their specific anti-*P. falciparum* IgG contents into high (>60 µg mL^-1^) or low (<60 µg mL^-1^) IgG. A list of the 38 samples used is shown in Table S2. Most samples used for further analysis belonged to the most frequent HLA-DRB1 genotype.

Immunoreactive epitopes in the five selected antigenic proteins were mapped by high density peptide microarrays designed to contain 15-mer amino acid sequence overlapped by the following 14 residues in the next 15-mer peptide to cover the whole sequence of the selected antigens. The reactivity of IgG antibodies present in 38 serum samples from the malaria endemic region was individually tested in the peptide microarrays. The IgG antigenicity profile of the five antigens along their sequence by these sera is shown in Figure 1. It is noteworthy that all five antigens showed a wide distribution of regions of IgG recognition along the protein sequences. The plots shown similar recognition patterns in both ASC and ANP samples. However, the overall signal was proportionally smaller in sera with low IgG values, suggesting that the ratio between the specific fraction of anti *P. falciparum* IgG and the total IgG remains constant among the population studied. To identify peptides that produce consistent reactivity among serum samples, a signal cut was established in the third quartile of the intensities recorded within each serum group. Figure 2 shows the frequency of serum samples emitting a fluorescent signal above the threshold in both ASC and ANP groups at all 15-mer peptide positions in the five antigens. According to the aggregated pattern of the individual sera, the peptide recognition frequencies are distributed similarly throughout the sequence, showing an almost complete coincidence between the ASC and ANP samples, which did not allow for the identification of any specific epitope of the subclinical state.

**Figure 1.**
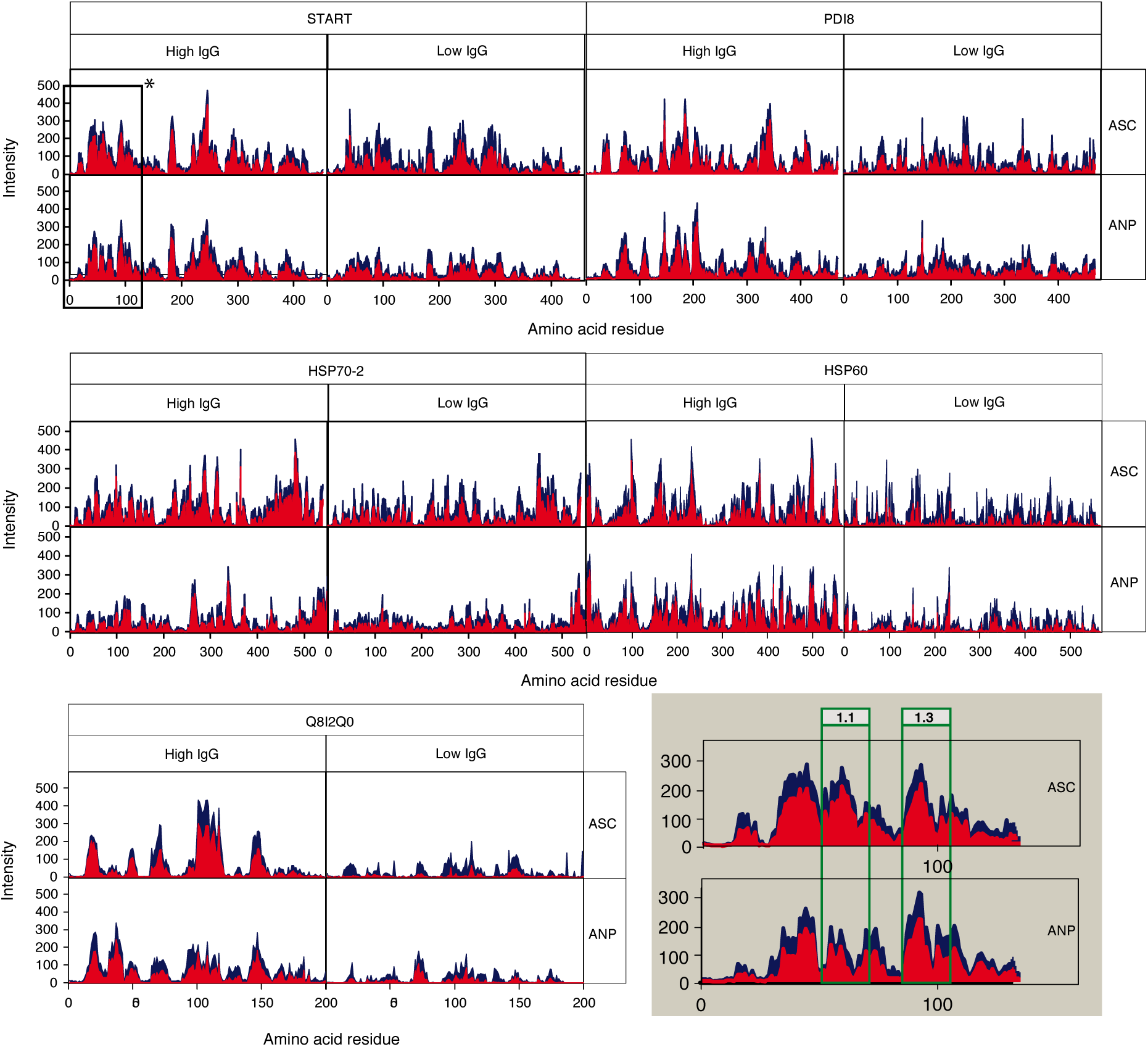
Epitope mapping in the five *P. falciparum* antigens. Data was recorded from 15-mer microarrays of overlapping peptides covering the whole protein sequence. Data shows average fluorescence *m* (blue) and standard error (*s*) subtracted average fluorescence *m-s* (red) along the sequence of the indicated antigens. Data is presented in a set of 4 plots to distinguish incubation with sera from ASC or ANP patients and for high IgG (> 60 μg mL^-1^) and low IgG sera. n values are as follow: 10 for ASC/High IgG; 8 for ANP/Low IgG; 11 for ASC/Low IgG; 9 for ANP/High IgG. * In the lower right corner (shaded box) is shown a magnification of microarrays immunoreactivity with high IgG sera in the START region corresponding to the position of the 20-mer selected peptides 1.1 and 1.3.

**Figure 2.**
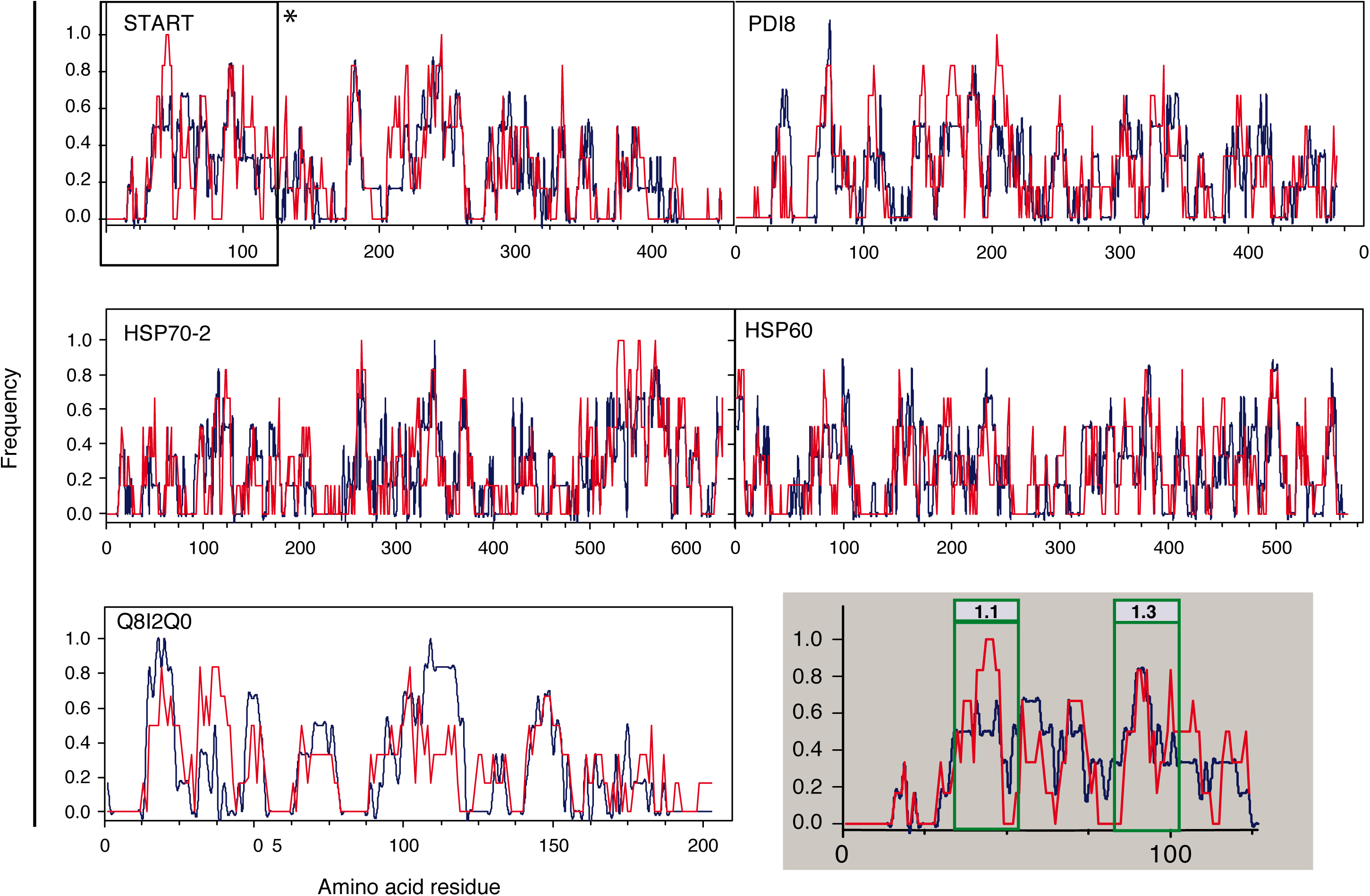
Comparison of IgG reactivity frequency between serum from adult patients with (blue: ASC) and without (red: ANP) parasitemia from Breman-Asikuma. X-axis shows amino acid sequence length of each antigen. Y-axis shows fluorescence signals above a cutoff stablished as the third quartile of the recorded data. Data from high IgG sera are shown. *Shadowed box at bottom right: enlargement of START region showing the position of the 20-mer selected peptides 1.1 and 1.3.

### Validation of highly immunoreactive epitopes

At the peak IgG signal along the antigen sequences, eight 20-mer length peptides were chosen that also showed the highest frequency of recognition by individual sera. Examples of the regions selected for START protein are depicted in Figures 1 and 2. The sequence of the chosen peptides and their location in the antigens are shown in Table 2. These eight synthesized peptides were tested for IgG reactivity using serum samples from the endemic malaria region. Figure 3 shows the amount of specific IgG for each 20-mer peptide detected in 8 ASC sera, 8 ANP sera and 4 control sera of individuals never exposed to the parasite. Although the signals recorded for the ANP group appear to be consistently higher than those found in the ASC group, no statistically significant differences could be determined for any of the peptides (P<0.05). When immunoreactivity was compared among ASC, ANP and control samples, only START and PDI8 peptides showed significant differences among them (see Figure 3). These results indicate that both START and PDI8 contain immunodominant epitopes that maintain high amounts of specific IgG in the sera of populations exposed to malaria. The constant immunoreactivity observed in sera from the ANP group that has not detectable parasites suggests a long persistence of this IgG after elimination of the infection.

**Table 2.**
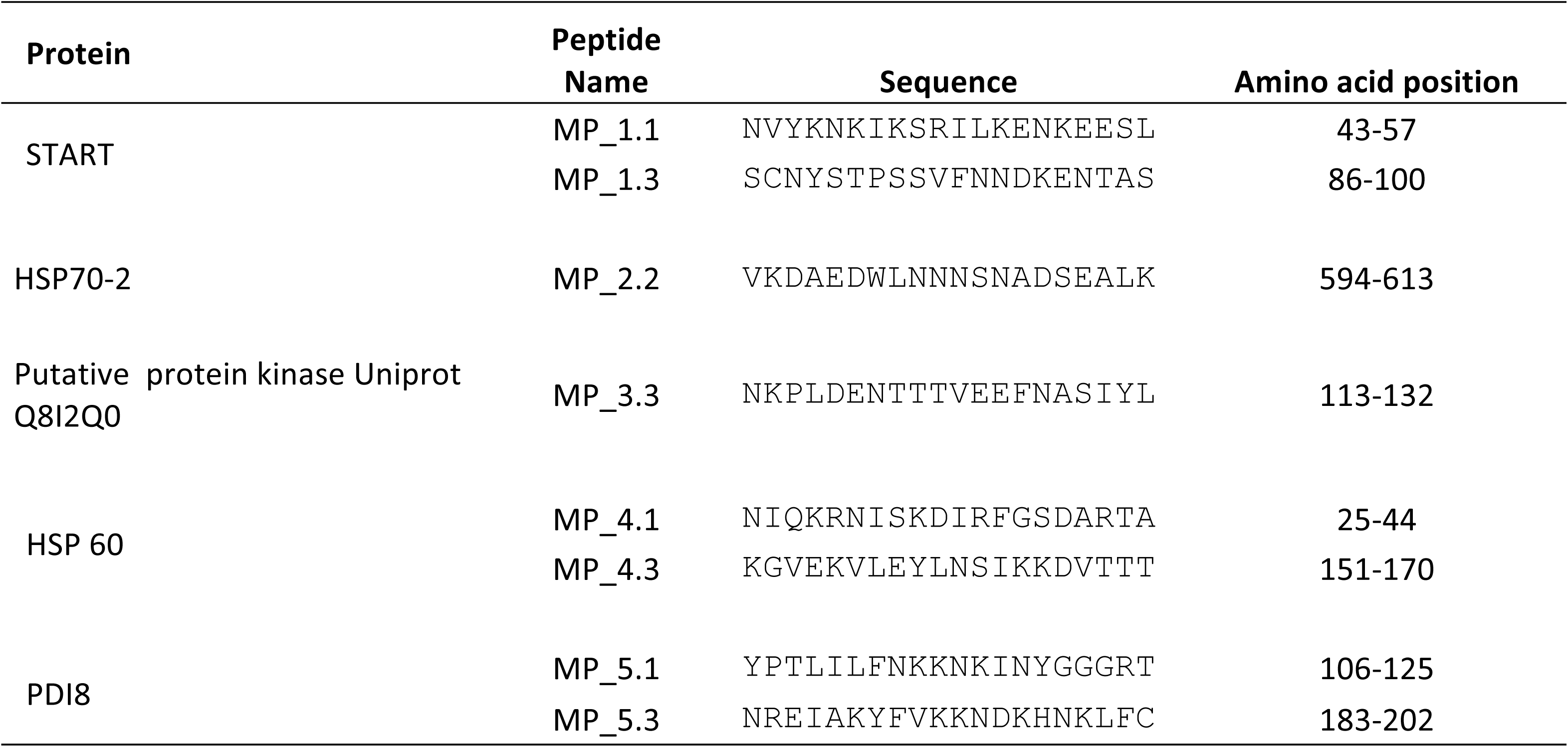
Peptides selected for IgG immunoreactivity analysis with patients sera from Breman-Asikuma (adults and children)

**Figure 3.**
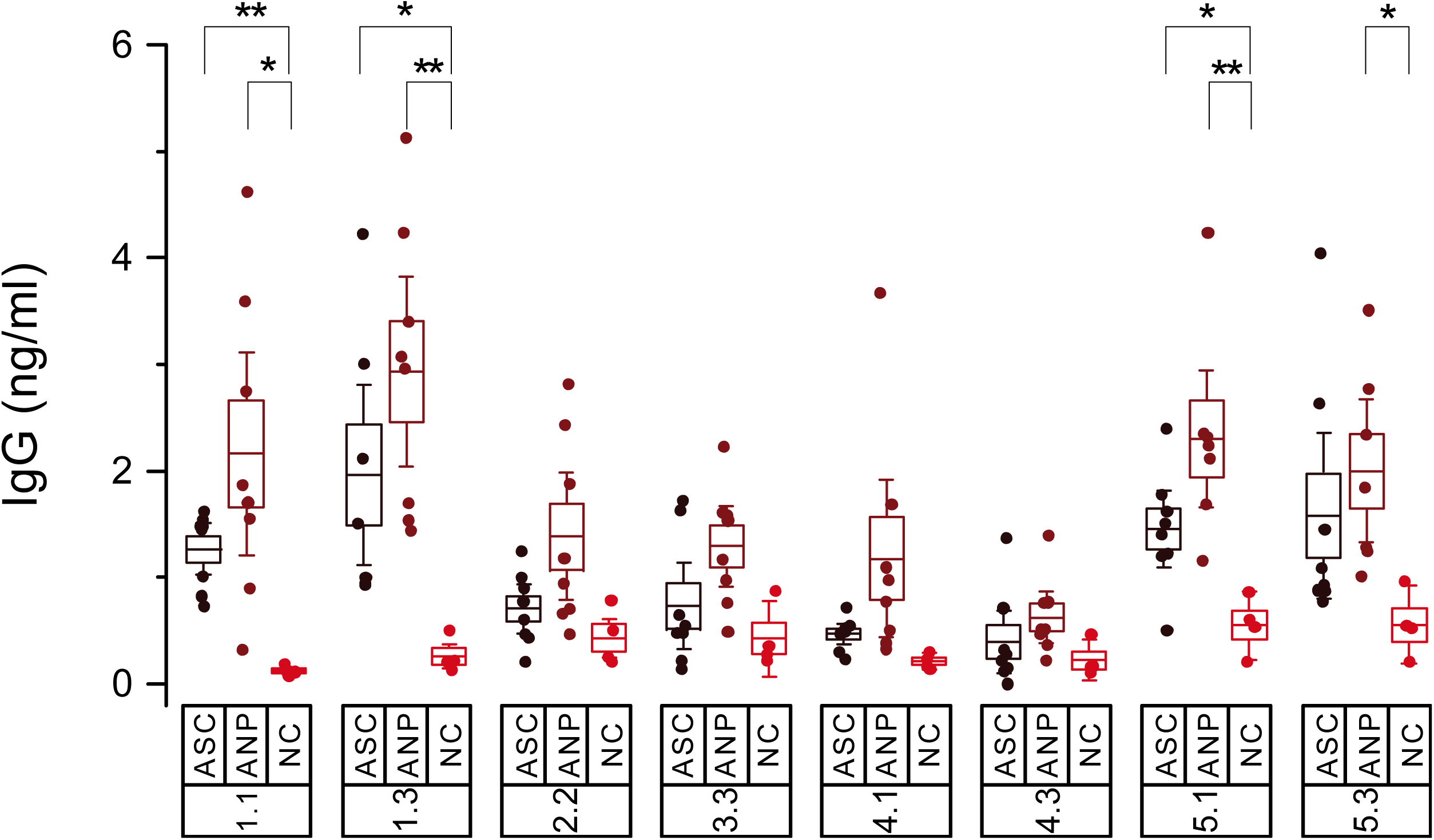
IgG reactivity of adult serum samples from the endemic malaria region with eight selected 20-mer epitopes. Data shows specific IgG concentration present in sera against 20-mer peptides sequences of START (1.1, 1.3); HSP70-2 (2.2); putative protein kinase Uniprot Q8I2Q0 (3.3); HSP60 (4.1, 4.3); and PDI8 (5.1, 5.3). Three different sera groups were used: asymptomatic subclinical (ASC, n=8), undetected malaria parasite (ANP, n=8) and no malaria exposed controls (NC, n=4). The 16 sera used from endemic malaria region were selected from the patients indicated with an asterisk in Table S2. Mean values are given by horizontal lines, covered with a box depicting standard error. Whiskers represent 90% confidence interval. T-test probability for significantly different means between groups are shown as: * P<0.05; ** P<0.01.

Since the persistence of anti-START and anti-PDI8 IgG in the adult population of the malaria-endemic region may be a consequence of repeated exposure to the parasite, it was conceivable that a less exposed population group would show a different pattern. To test this hypothesis, serum samples from children younger than 5 years old in the same endemic area were analyzed to detect specific IgG against peptides 1.1, 1.3, 5.1 and 5.3 that were the most immunoreactive of START and PDI-8 in adults. In this trial, the sera of children with symptoms of malaria or asymptomatic, corresponding to those showing parasites by microscopic screening (parasitemia >0.1 %, clinical malaria: CCM), those for whom *P. falciparum* was detected only by PCR of its DNA but not by microscopy (subclinical malaria: CSC), and those who did not show the presence of the parasite by either method (CNP). The results, shown in Figure 4, confirm the immunoreactivity of both antigens in sera collected from children, but unlike adults, the signal obtained in cases of subclinical malaria was significantly higher than the signal in uninfected cases (p<0.05).

**Figure 4.**
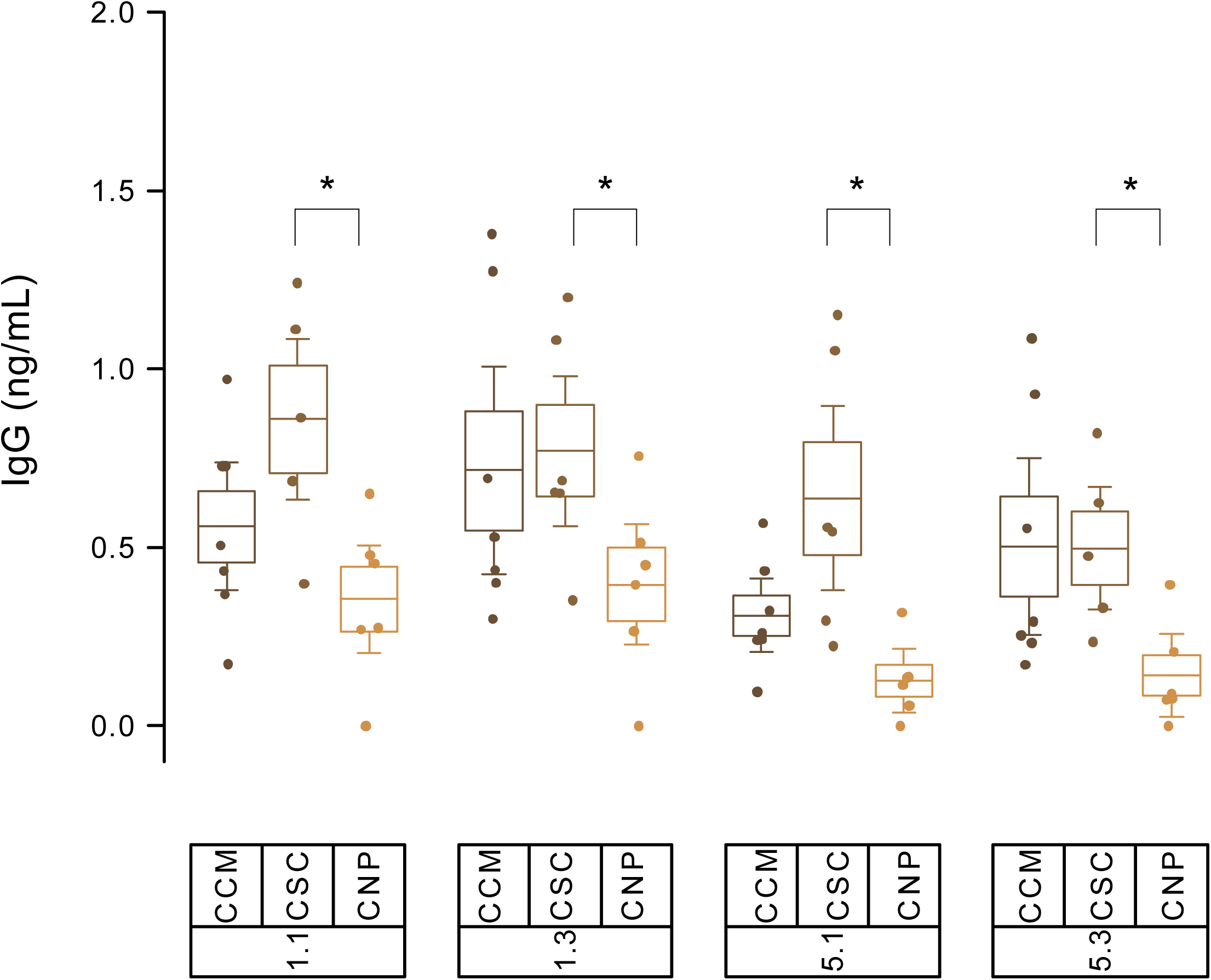
IgG reactivity of children serum samples from the endemic malaria region with four selected 20-mer epitopes. Data shows specific IgG concentration present in sera against 20-mer peptides sequences of START (1.1, 1.3) and PDI8 (5.1, 5.3). Three different sera groups were used: clinical malaria samples (CCM, parasitemia >0.1%); asymptomatic subclinical (CSC), and undetected malaria parasite (CNP). Mean values are given by horizontal lines, covered with a box depicting standard error. Whiskers represent 90% confidence interval. T-test probability for significantly different means between groups are shown as: * P<0.05; ** P<0.01.

## Discussion

Despite the variety of *P. falciparum* antigenic proteins identified by different approaches (10, 11, 32-35), only a few have been shown to be; albeit partially, relevant to malaria immunity (10) or for detection purposes (36, 37). Previously, the humoral response to *P. falciparum* in patients with imported malaria was used to identify immunogenic proteins by western blot 1D analysis. (32). Here, we have used a 2D immunoproteomics approach using imported malaria patient sera to increase search resolution, mapped the position of potential epitopes in a selection of five immunodominant antigens, and validated their immunogenicity using serum samples from individuals in a malaria-endemic region. Although imported malaria samples used for immunoproteomics were classified into two groups based on different parasitemias, a clear association between individual antigens and disease stage could not be determined. In fact, no significant differences in total serum anti-*P. falciparum* IgG levels were found between the two groups. However, a higher overall avidity for anti-*P. falciparum* IgG was recorded for the serum group of patients with the lower parasitic load (submicroscopic). Recently it has been shown that the antibody avidity along the course of infection for two antigens that are targets of naturally acquired immunity EBA175 (region RIII-V) and *Pf* Reticulocyte Homologue Protein 2 (*Pf*Rh2) is variable between individuals, showing in some cases the typical increase in affinity due to maturation of the B cell or, in other cases, a decrease over time (38). Therefore, the overall decrease in anti-*P. falciparum* IgG avidity observed in the imported malaria group with clinical symptoms (patients with diminished exposure to the parasite and high parasitemia) can be explained by progressive loss of affinity in the main specific antibodies or by replacement of high affinity IgG by a different subset targeting different antigens, or by the association of both effects. Thus, greater avidity could be considered as a protective factor against infection, as individuals with more avid IgG have less parasitemia values as observed upon vaccination with RTS,S/AS01E (39). Eight of the 17 antigens identified by immunoproteomics had not been previously reported, including experimental approaches and computational prediction of the entire proteome (10). This supports the notion that broad and diverse experimental approaches are necessary to obtain a more complete range of *P. falciparum* immunogenic proteins. Chaperones were the most frequently occurring proteins in the fate category, including the heat shock proteins HSP70, HSP70-2, HSP70-3 and HSP60. In the metabolic category, Plasmepsins I and II, involved in hemoglobin digestion, were similarly found in sera from both imported malaria groups. Plasmepsin I has been detected in intraerythrocytic young stages whereas Plasmepsin II expression has been associated with mature stages (40). A similar distribution was observed in the generation of antibodies against dihydrolipoyl dehydrogenase and dihydrolipoamide acyltransferase, both of them components of the multienzyme pyruvate dehydrogenase complex. Two further metabolism-related proteins from the parasite, ferrodoxin reductase-like protein, involved in fatty acid metabolism, and vacuolar ATP synthase subunit b, likely involved in acidification of the digestive vacuole (41), were also found in sera from both groups. However, while antibodies against ferrodoxin-reductase-like protein were found mainly in patients with clinical symptoms, the vacuolar ATP synthase subunit b was more antigenic in those individuals without clinical signs of malaria. The most frequently detected antigen was initially identified as a conserved protein of unknown function Uniprot Q8I2Q0, corresponding to gene *PF*3D7_0925900 in the genome of *P. falciparum* 3D7. Further analysis by sequence comparison (not shown) revealed that this sequence may correspond to an uncomplete gene duplication of a putative protein kinase Uniprot Q8I2P9, corresponding to gene *PF*IT_*PF*I1270W, located 20 kpb downstream *PF*3D7_0925900. Within the cell surface class, three immunogenic proteins which assist in the process of erythrocyte invasion (42) were identified. Two of them, MSP-1 and MSP-9, are located at the merozoite surface (43) and RAP2, associated with the low-molecular-weight rhoptry complex in the rhoptry organelles (44). Finally, a new antigenic protein, integral membrane STAR-related lipid transport START, was found in about 55% of the samples from all three groups. Although the humoral response against some of these proteins has previously been reported in other studies using samples from malaria-endemic regions (11, 33-35) others such as putative protein kinase, HSP70-2, V-Type proton ATPase subunit B, START, LPD1, PDI8, ferredoxin reductase-like protein and plasmepsins I and II, they may be indicative of differences in the specific IgG range shown in individuals with imported malaria caused only by an isolated infection in a partially protected individual.

To investigate epitope immunoreactivity in the identified antigens, five of them were selected for mapping of their epitopes by humoral response with sera from an endemic malaria population. Putative protein kinase Q8I2Q0, HSP70-2, HSP60 and START were the most frequently detected antigens in ISC sera, suggesting that these proteins may induce a strong humoral response that lasts after parasite clearance. These five selected proteins are not only present in the asexual stage of merozoite (29) guaranteeing potential immunogenic contact by the host during the intraerythrocytic cycle (see Table 1), but have also recently been guaranteed gene expression throughout the asexual and sexual stages of the Plasmodium cycle at discrete levels (45), thus implying non-saturating exposure to the immune system. START antigen was also detected even more frequently than other well-known antigenic membrane proteins, such as MSPs. The protein disulfide isomerase PDI8 was also included in the epitope mapping as it was also detected with high frequency and its function associated with multiple proteins during the parasite cycle suggests that it may be widely exposed to the host immune system. Complete mapping of potential linear epitopes in these proteins using overlapping 15-mer peptide microarrays revealed high IgG reactivity sequences homogeneously distributed throughout the sequence in all antigens analyzed. As the collection of sera used for antigen screening was confined to asymptomatic adults, potential epitopes were expected to represent sequences associated with acquired immunity and/or subclinical infections. Subsequent ELISA analysis of a selection of 20-mer peptides showing both a high IgG recognition signal and a high binding frequency in endemic malaria samples allowed quantification of specific IgG concentrations in sera. Since no statistically significant differences in recognition in microarray patterns or in the IgG binding levels to the 20-mer peptides were observed between ASC endemic region samples with subclinical infection and non-infected ANP controls, we can conclude that the host IgG collection directed against these epitopes represents a subpopulation of long-term IgG antibodies present in individuals after repeated exposure to the parasite.

The persistence of anti-malarial IgG response may be due to a long-lasting B-cell memory or a high number of long-lasting plasma cells (LLPC) (46). On the other hand, the longevity of the antibody response observed in the adult population of malaria-endemic regions has been explained as a result of a progressive increase in LLPC over the course of recurrent reinfections by highly antigenic variable parasites (47). The titration of specific IgG bound to different epitopes of the five selected antigens revealed that only START and PDI8 epitopes yielded a significantly higher IgG signal above background in the malaria-endemic sera. Humoral response against START and PDI8 in uninfected individuals from endemic areas may be caused by repeated exposure to the parasite and increased production of specific IgG against these two proteins by LLPC.

To further explore this possibility, serum samples from children under 5 years old from the same endemic area were analyzed. It is now accepted that resistance to severe malaria is rapidly attained within the first five years of life (48), while immunity to non-severe malaria is slowly acquired over years of repeated re-infection, often leading to a subclinical course of the disease that can only be detected by PCR amplification (49). Remarkably, only *P. falciparum*-infected children displayed anti-START or anti-PDI8 IgGs, indicating that the presence of these antibodies in adults may be associated with the development of acquired immunity against non-severe malaria. Taken together, these results suggest that *P. falciparum* antigens involved in the maintenance of acquired immunity show a wide range of B-cell epitopes of which at least the START and PDI8 epitopes lead to the production of a long-lasting IgG response that is maintained after parasite clearance, which may be indicative of a relevant role of these antigens in naturally acquired immunity. In addition, the presence of anti-START and/or anti-PDI8 IgG in children with subclinical malaria would also be useful as a marker for such undetected reservoir infection. Thus, these epitopes could be used as a rapid method of ELISA immunodetection at large-scale in elimination programs instead of PCR diagnosis.

## Materials and Methods

### Sample collection and population

Four sets of human sera were used in the present study. The first set (n=19) was obtained from adult sub-Sahara African patients resident in Spain with imported malaria detected after a recent trip from Africa (Table S1). The second set (n=38) was obtained from adult donors (17-40 years old, excluding pregnant women) living at Breman-Asikuma (Ghana) attending to a local hospital; Our Lady of Grace Hospital, but showing no symptoms of malaria or other endemic infections (tuberculosis, hepatitis B, hepatitis C, HTLV, dengue and HIV) at the time of collection (Table S2I). The third set (n=20) was collected at the same location from children aged 6 weeks to 5 years old, including patients showing malaria symptoms (Table S3).

A fourth set of 4 European volunteers never exposed to malaria was used as a control. This study was approved by the corresponding Ethical Review Boards at Research Institute Hospital 12 de Octubre, Madrid and the University of Ghana’s College of Health Sciences.

### Protein extracts from in vitro cultures of *Plasmodium falciparum*

Clinical *P. falciparum* isolate UCM7 (28) from West Africa was cultured to high parasitemia (> 50%) following the previously described method (50). Briefly, the culture media consisted of standard RPMI 1640 (Sigma-Aldrich) supplemented with 0.5% Albumax I (Gibco), 100 µM Hypoxanthine (Sigma-Aldrich), 25 mM 101 HEPES (Sigma-Aldrich), 12.5 µg/mL Gentamicine (Sigma-Aldrich) and 25 mM NaHCO_3_ (Sigma-Aldrich). Culture at 37°C with 5% CO_2_ started by mixing a cryopreserved stock with fresh erythrocytes to achieve 1% hematocrit. Growth was monitored by microscopy in thin blood smears and media changes and synchronization steps following defined criteria (50) according to parasite stage and parasitemia. Protein extracts from *P. falciparum* isolate UCM7 were obtained from three 150 ml culture flasks of infected red blood cells (iRBC) synchronized to mature forms (schizonts) at high parasitemia (50). Cultures were harvested and erythrocytes resuspended 1:1 in PBS containing 0.1% saponin to gently vortex during 15s for lysing RBC. Released parasites were centrifuged at 800xg during 5 min and the pellet was 3x washed in cold PBS to subsequently centrifuged at 15,000xg, 10 min. Pellets were solubilized in 50 mM Tris-HCl pH8 containing 50mM NaCl, 3% CHAPS, 0.5% MEGA 10 and protease inhibitor cocktail from Roche. Resuspension was gently mixed at 4°C during 15 min followed by 4x freeze-thaw cycles. Cold (4°C) centrifugation at 15,000xg during 10 min resulted in a supernatant considered the parasite extract. Protein concentration was quantified using Bradford based *Protein Assay* from Bio-Rad and total protein lysates stored at −20°C until use.

### Total and *P. falciparum* specific immunoglobin-G quantification

Immunoglobin-G (IgG) in sera samples was quantified using indirect IgG ELISA detection kit (Bethyl Laboratories) following the manufacturer’s instructions. For total IgG quantification, microtiter plates were coated with 1μg/well of anti-human IgG (Bethyl Laboratories) in 100μl carbonate/bicarbonate buffered solution (Sigma C3041) for 1 hour at room temperature; while specific anti-*P. falciparum* IgG were quantified using 0.5 μg/well of parasite protein extract (see above) in the same buffer as coating antigen overnight at 4°C. From this step onwards, all wells followed same protocol. Plates were blocked with 1% bovine serum albumin (BSA) in Tris-buffered saline solution and diluted human serum was incubated for 1 h at room temperature, using 1:50,000 dilution for total IgG and 1:5,000 dilution for specific IgG quantification. IgG binding was detected with goat anti-human IgG conjugated with horseradish peroxidase (HRP) at 1:75,000 dilution. The enzymatic reaction was developed using 3,3’,5,5’ tetramethyl benzidine (TBM) as enzyme substrate and stopped by the addition of 100 µl of 4 N H_2_SO_4_ Absorbance readings were obtained at 652 nm in a Varian Cary 50 Biospectrophotometer. Serum samples from non-exposed European individuals were used to establish the negative cut-off of the ELISA (mean OD+2 S.D.). Human reference serum from Bethyl Laboratories was used to generate a logistic four-parameter sigmoidal standard curve.

Antibody avidity determination of sera from imported malaria patients is given in Supplementary Information.

### Immunoproteomics for antigen identification

*P. falciparum* UCM7 extracts were run in 2D-PAGE (300 µg for mass-spectrometry identification / 150 µg for immunoblot) for subsequent Western blot analysis using sera from patients with imported malaria. Protein extract were cup loaded onto IPG strips (pH 3−11, 18 cm) previously hydrated overnight with a buffer containing 8 M urea, 2 M thiourea, 4% (w/v) CHAPS, 1.2%, DeStreak reagent and 2% ampholites pH 3-11. For first-dimensional separation, IEF was performed using the IPGphor 3 IEF system (GE Healthcare) at 20 °C. Voltage was gradually ramped in a step-and-hold manner to 1,000 V in three steps: 1 h at 120 V, 1 h at 500 V, and 2 h at 500−1,000 V. Then, voltage increased to 4,000 V in linear gradient (1,000−4,000 V) over the next 9 hours. Run terminated after ≈ 70,000 V/h. The focused strips were equilibrated twice with gentle vertical shaking. The first equilibration during 12 min was performed with a solution containing 100 mM Tris-HCl pH 8.8, 6 M urea, 30% glycerol, 2% SDS and 0.5% DTT. In the second equilibration solution, DTT was replaced with 4.5% iodoacetamide and incubation continued 5 min. SDS-PAGE using a Ettan-Daltsix unit at 2 W/gel for 30 min followed by 20 W/gel for 4 h was run on homogeneous 12.5% pre-casted gels. To visualize and further excise and identify protein spots, the 2D-gels loaded with 300 µg were fixed in 10% methanol/7% acetic acid for 30 min and overnight stained with Sypro Ruby (Sigma). Two 2D-gels for staining and protein-spot excising (300 µg) and 20 replicated 2D-gels for Western-blot (150 µg) were run in parallel, in group of 6, under identical voltage conditions to warrant replica. The 2D-gels loaded with 150 µg were transferred to polyvinylidene difluoride (PVDF) membranes. Transfer efficiency was confirmed by: (i) gel staining with Sypro Tangerine before transfer; and (ii) membrane staining with Sypro Ruby after transfer. Stained gels or blots were scanned in a Typhoon 9400 variable mode imager (GE Healthcare). For immunodetection with individual patient serum, the transferred PVDF membranes were blocked for 2 hours in 10% non-fat skimmed milk (w/v) in PBS containing 0.05% Tween-20 followed by overnight incubation with patient serum at 1: 10,000 dilution. Then, goat anti-human IgG HRP linked (Amersham Bioscience) as secondary antibody was incubated for 1 h at room temperature at a 1:10,000 dilution. Antigen-antibody reaction was visualized using the SuperSignal chemiluminescent substrate (Pierce) and exposure to X-ray film. Human sera from Europeans never exposed to malaria were used as negative controls. Gel and immunoblot images were analyzed using PDQuest software, version 8.0.1 (BioRad Laboratories Inc., Munich, Germany), and spots of interest were manually excised from the 2D gels, in-gel reduced, alkylated, and then digested with trypsin as described elsewhere (51). Excised spots were washed twice in water, shrunk for 15 min with 100% acetonitrile, and dried in a Savant SpeedVac for 30 min. The samples were then reduced using 10 mM DTT in 25 mM ammonium bicarbonate for 30 min at 56°C and subsequently alkylated with 55 mM iodoacetamide in 25 mM ammonium bicarbonate for 20 min in the dark. Finally, samples were digested with 12.5 ng/µL sequencing grade trypsin in 25 mM ammonium bicarbonate (pH 8.5) overnight at 37 °C. After digestion, the supernatant was collected, and 1 µL was spotted onto a MALDI target plate and allowed to air-dry at room temperature. Next, 0.4 µL of a 3 mg/mL of α-cyano-4-hydroxytranscinnamic acid matrix in 50% acetonitrile was added to the dried peptide spots and allowed, again, to air-dry at room temperature. MALDI time-of-flight (TOF) MS analyses were performed in a 4800 Proteomics Analyzer MALDI-TOF/TOF mass spectrometer (Applied Biosystems, Warrington, U.K.) operated in positive reflector mode, with an accelerating voltage of 20,000 V. Mass spectra were then acquired for peptide mass fingerprinting (PMF). Proteins were identified by comparing the trypsin-digested peptide masses with the data provided in two databases (NCBI) residues and UniprotKB-SwissProt v.56.6) separately using MASCOT 1.9 (http://www.matrixscience.com) through the Global Protein Server v3.5 from Applied Biosystems. All of the identified proteins fulfilled the criterion of being significant (*p* < 0.005) according to probability based on the Mowse score (52). Statistical analyses were performed using the Statgraphics Plus 5.1 software package.

### High-density Peptide Arrays

Sera samples from subjects from Breman-Asikuma were tested for the recognition of peptides in arrays. They were synthesized on glass slides using photolithographic peptide array (53) using a layout with 12 identical sectors per slide. Each sector contained 6630 peptide fields. The surface coating on the slides was made by incubation of Nexterion E slides (Schott AG) with a 2% w/v linear copolymer of N,N’-dimethylacrylamide and aminoethyl methacrylate as described (54). Each sector of the arrays was incubated with a different serum diluted 1:200 in buffer (0.05 M Tris/acetate pH 8.0, 0.13 M NaCl, 0.1% v/v Tween20, 1g/L BSA) for 2 hours in after blocking. Then, after washing in same buffer, the slides were incubated for 2 hours at room temperature with 1 µg/mL Cy3-conjugated goat anti-human IgG (H+L) (Sigma) and washed again. The slides were centrifuged in a slide spin-dryer and scanned at 1 µm resolution in an InnopSys 900 laser scanner using an excitation wavelength of 532 nm. The images were analyzed using the PepArray analysis program (Schafer-N) and the fluorescent signal in each field was calculated as the weighted average of the pixel intensities in that field.

### Peptide-based ELISA

Specific IgG binding in serum samples to selected peptides was tested by ELISA. Microtiter plates (Maxisorb, NUNC) were coated in duplicate with peptides by adding 100 µL of a 2.5 µg mL^-1^ solution in 0.05 M Carbonate-Bicarbonate buffer, pH 9.6. After overnight incubation at 4°C, plates were washed with 50 mM Tris, 0.14 M NaCl, 0.05% Tween 20, pH 8.0 buffer five times, 200 µL of blocking buffer (50 mM Tris, 0.14 M NaCl, 1% BSA, pH 8.0) were added and incubated at room temperature for 30 minutes. After one more wash, 100 µL of serum diluted 1:450 in wash buffer was added. After incubation at room temperature during 60 min, the supernatant was removed, the wells were washed once more and 100 µL of goat Anti-IgG-Fc HRP conjugated at 1 mg mL^-1^ (Bethyl) were added at 1:15000 dilution. HRP activity was detected as indicated above.

## Acknowledgements

We thank all the patients and parents who participated in this study for their contribution. We also thank to Maria L. Ruperez, Edwige Gaba and Juana Garrido and all the Breman-Asikuma Community of the Sisters of Charity of St. Anne for strategic support and logistics and Emmanuel Sekyere and all the laboratory staff at Our Lady of Grace Hospital for excellent technical assistance. We also thank students Alejandra Martínez-Serna, Luis Diez Espallargas, Ana Abad and Sara Marcos for data collection. This work was supported by Spanish-MINECO grants BIO2013-44565R and BIO2016-77430R and a research fellowship to P.A. from Universidad Complutense de Madrid.

## References

1. WHO (2016) WHO World Malaria Report 2016. (WHO, Geneva).

2. Murray CJ, et al. (2012) Global malaria mortality between 1980 and 2010: a systematic analysis. Lancet 379(9814):413–431.

3. Umbers AJ, Aitken EH, & Rogerson SJ (2011) Malaria in pregnancy: small babies, big problem. Trends Parasitol 27(4):168–175.

4. Monge-Maillo B & Lopez-Velez R (2012) Migration and malaria in europe. Mediterr J Hematol Infect Dis 4(1):e2012014.

5. McGregor IA (1974) MECHANISMS OF ACQUIRED IMMUNITY AND EPIDEMIOLOGICAL PATTERNS OF ANTIBODY-RESPONSES IN MALARIA IN MAN. Bulletin of the World Health Organization 50(3-4):259–266.

6. Langhorne J, Ndungu FM, Sponaas A-M, & Marsh K (2008) Immunity to malaria: more questions than answers. Nature Immunology 9(7):725–732.

7. Cohen S, Carrington S, & McGregor IA (1961) Gamma-globulin and acquired immunity to human malaria. Nature 192(480):p733-&.

8. Healer J, Chiu CY, & Hansen DS (2018) Mechanisms of naturally acquired immunity to P-falciparum and approaches to identify merozoite antigen targets. Parasitology 145(7):839–847.

9. Sercarz EE, et al. (1993) Dominance and crypticity of t-cell antigenic determinants. Annual Review of Immunology 11:729–766.

10. Doolan DL, et al. (2003) Identification of Plasmodium falciparum antigens by antigenic analysis of genomic and proteomic data. P Natl Acad Sci USA 100(17):9952–9957.

11. Crompton PD, et al. (2010) A prospective analysis of the Ab response to Plasmodium falciparum before and after a malaria season by protein microarray. Proc Natl Acad Sci U S A 107(15):6958–6963.

12. Yewdell JW (2006) Confronting complexity: Real-world immunodominance in antiviral CD8(+) T cell responses. Immunity 25(4):533–543.

13. Sela-Culang I, Kunik V, & Ofran Y (2013) The structural basis of antibody-antigen recognition. Frontiers in Immunology 4.

14. Van Regenmortel MHV (2006) Immunoinformatics may lead to a reappraisal of the nature of B cell epitopes and of the feasibility of synthetic peptide vaccines. Journal of Molecular Recognition 19(3):183–187.

15. Dudek NL, Perlmutter P, Aguilar M-I, Croft NP, & Purcell AW (2010) Epitope Discovery and Their Use in Peptide Based Vaccines. Current Pharmaceutical Design 16(28):3149–3157.

16. Sanchez-Trincado JL, Gomez-Perosanz M, & Reche PA (2017) Fundamentals and Methods for T- and B-Cell Epitope Prediction. J Immunol Res 2017:2680160.

17. Gomara MJ & Haro I (2007) Synthetic peptides for the immunodiagnosis of human diseases. Current Medicinal Chemistry 14(5):531–546.

18. Vekemans J & Ballou WR (2008) Plasmodium falciparum malaria vaccines in development. Expert Review of Vaccines 7(2):223–240.

19. Atassi MZ & Smith JA (1978) PROPOSAL FOR NOMENCLATURE OF ANTIGENIC SITES IN PEPTIDES AND PROTEINS. Immunochemistry 15(8):609–610.

20. Rubinstein ND, et al. (2008) Computational characterization of B-cell epitopes. Molecular Immunology 45(12):3477–3489.

21. Balam S, et al. (2014) Plasmodium falciparum merozoite surface protein 2: epitope mapping and fine specificity of human antibody response against non-polymorphic domains. Malaria J 13.

22. Epping RJ, et al. (1988) AN EPITOPE RECOGNIZED BY INHIBITORY MONOCLONAL-ANTIBODIES THAT REACT WITH A 51 KILODALTON MEROZOITE SURFACE-ANTIGEN IN PLASMODIUM-FALCIPARUM. Molecular and Biochemical Parasitology 28(1):1–10.

23. Rodrigues-da-Silva RN, et al. (2016) In silico Identification and Validation of a Linear and Naturally Immunogenic B-Cell Epitope of the Plasmodium vivax Malaria Vaccine Candidate Merozoite Surface Protein-9. Plos One 11(1).

24. Chougnet C, et al. (1991) LYMPHOPROLIFERATIVE RESPONSES TO SYNTHETIC PEPTIDES FROM MEROZOITE RING-INFECTED ERYTHROCYTE SURFACE-ANTIGEN AND CIRCUMSPOROZOITE PROTEIN - A LONGITUDINAL-STUDY DURING A FALCIPARUM-MALARIA EPISODE. American Journal of Tropical Medicine and Hygiene 45(5):560–566.

25. Avila SLM, et al. (2001) Immune responses to multiple antigen peptides containing T and B epitopes from Plasmodium falciparum circumsporozoite protein of Brazilian individuals naturally exposed to malaria. Parasite Immunology 23(2):103–108.

26. Troyeblomberg M, et al. (1989) T-CELL AND B-CELL RESPONSES OF PLASMODIUM-FALCIPARUM MALARIA-IMMUNE INDIVIDUALS TO SYNTHETIC PEPTIDES CORRESPONDING TO SEQUENCES IN DIFFERENT REGIONS OF THE P-FALCIPARUM ANTIGEN PF155 RESA. Journal of Immunology 143(9):3043–3048.

27. Burbelo PD, Ching KH, Bush ER, Han BL, & Iadarola MJ (2010) Antibody-profiling technologies for studying humoral responses to infectious agents. Expert Review of Vaccines 9(6):567–578.

28. Linares M, et al. (2011) Malaria hidden in a patient with diffuse large-B-cell lymphoma and sickle-cell trait. J Clin Microbiol 49(12):4401–4404.

29. Florens L, et al. (2002) A proteomic view of the Plasmodium falciparum life cycle. Nature 419(6906):520–526.

30. Alonso-Padilla J, Lafuente EM, & Reche PA (2017) Computer-Aided Design of an Epitope-Based Vaccine against Epstein-Barr Virus. J Immunol Res 2017:9363750.

31. Quinzo MJ, Lafuente EM, Zuluaga P, Flower DR, & Reche P A, (2019) Computational assembly of a human Cytomegalovirus vaccine upon experimental epitope legacy. BMC Bioinformatics.

32. Costa RM, Nogueira F, de Sousa KP, Vitorino R, & Silva MS (2013) Immunoproteomic analysis of Plasmodium falciparum antigens using sera from patients with clinical history of imported malaria. Malaria J 12.

33. Trieu A, et al. (2011) Sterile protective immunity to malaria is associated with a panel of novel P. falciparum antigens. Mol Cell Proteomics 10(9):M111 007948.

34. Baum E, et al. (2013) Protein Microarray Analysis of Antibody Responses to Plasmodium falciparum in Western Kenyan Highland Sites with Differing Transmission Levels. Plos One 8(12).

35. Uplekar S, et al. (2017) Characterizing Antibody Responses to Plasmodium vivax and Plasmodium falciparum Antigens in India Using Genome-Scale Protein Microarrays. Plos Neglected Tropical Diseases 11(1).

36. Druilhe P, Moreno A, Blanc C, Brasseur PH, & Jacquier P (2001) A colorimetric in vitro drug sensitivity assay for Plasmodium falciparum based on a highly sensitive double-site lactate dehydrogenase antigen-capture enzyme-linked immunosorbent assay. American Journal of Tropical Medicine and Hygiene 64(5-6):233–241.

37. Tritten L, Matile H, Brun R, & Wittlin S (2009) A new double-antibody sandwich ELISA targeting Plasmodium falciparum aldolase to evaluate anti-malarial drug sensitivity. Malaria J 8.

38. Tijani MK, et al. (2018) Factors influencing the induction of high affinity antibodies to Plasmodium falciparum merozoite antigens and how affinity changes over time. Scientific Reports 8.

39. Dobano C, et al. (2019) Concentration and avidity of antibodies to different circumsporozoite epitopes correlate with RTS, S/AS01E malaria vaccine efficacy. Nat Commun 10(1):2174.

40. Francis SE, Banerjee R, & Goldberg DE (1997) Biosynthesis and maturation of the malaria aspartic hemoglobinases plasmepsins I and II. Journal of Biological Chemistry 272(23):14961–14968.

41. Karcz SR, Herrmann VR, Trottein F, & Cowman AF (1994) CLONING AND CHARACTERIZATION OF THE VACUOLAR ATPASE-B SUBUNIT FROM PLASMODIUM-FALCIPARUM. Molecular and Biochemical Parasitology 65(1):123–133.

42. Cowman AF & Crabb BS (2006) Invasion of red blood cells by malaria parasites. Cell 124(4):755–766.

43. Gilson PR, et al. (2006) Identification and stoichiometry of glycosylphosphatidylinositol-anchored membrane proteins of the human malaria parasite Plasmodium falciparum. Mol Cell Proteomics 5(7):1286–1299.

44. Saul A, et al. (1992) The 42-kilodalton rhoptry-associated protein of *Plasmodium falciparum*. Molecular and Biochemical Parasitology 50(1):139–150.

45. Howick VM, et al. (2019) The Malaria Cell Atlas: Single parasite transcriptomes across the complete Plasmodium life cycle. Science 365(6455).

46. White MT, et al. (2014) Dynamics of the Antibody Response to Plasmodium falciparum Infection in African Children. Journal of Infectious Diseases 210(7):1115–1122.

47. Hviid L, Barfod L, & Fowkes FJI (2015) Trying to remember: immunological B cell memory to malaria. Trends in Parasitology 31(3):89–94.

48. Cowman AF, Healer J, Marapana D, & Marsh K (2016) Malaria: Biology and Disease. Cell 167(3):610–624.

49. Okell LC, Ghani AC, Lyons E, & Drakeley CJ (2009) Submicroscopic Infection in Plasmodium falciparum-Endemic Populations: A Systematic Review and Meta-Analysis. Journal of Infectious Diseases 200(10):1509–1517.

50. Radfar A, et al. (2009) Synchronous culture of Plasmodium falciparum at high parasitemia levels. Nat Protoc 4(12):1899–1915.

51. Shevchenko A, Tomas H, Havlis J, Olsen JV, & Mann M (2006) In-gel digestion for mass spectrometric characterization of proteins and proteomes. Nat Protoc 1(6):2856–2860.

52. Pappin DJC, Hojrup P, & Bleasby AJ (1993) RAPID IDENTIFICATION OF PROTEINS BY PEPTIDE-MASS FINGERPRINTING. Current Biology 3(6):327–332.

53. Buus S, et al. (2012) High-resolution Mapping of Linear Antibody Epitopes Using Ultrahigh-density Peptide Microarrays. Mol Cell Proteomics 11(12):1790–1800.

54. Hansen LB, Buus S, & Schafer-Nielsen C (2013) Identification and Mapping of Linear Antibody Epitopes in Human Serum Albumin Using High-Density Peptide Arrays. Plos One 8(7).

