## Supplemental text, figures and tables for "*Plasmodium falciparum* immunodominant IgG epitopes in subclinical malaria"

**This PDF file includes:**

Supplementary text
Figures S1 to S2
Tables S1 to S4
SI References

### SUPPLEMENTARY TEXT

#### Material and Methods for data shown in Supplementary Figure S1 and Supplementary Table S2.

#### Determination of IgG avidity (Figure S1)

To test antibody avidity within the serum samples from imported malaria patients, seven different concentrations (0, 0.5, 1, 2, 3, 4 and 6 M) of NaSCN were used to disrupt antigen-antibody binding along the IgG determination protocol (section 1.3 in Material and Methods) according to establish procedures (1). After the serum incubation step, wells were 3x washed using PBS containing 0.05% Tween-20. Next, each of the above NaSCN concentrations were added to a different well. Microplates were allowed to stand at room temperature for 15 min and extensively washed (x6) with PBS containing 0.05% Tween-20. Subsequent steps were performed as described in the IgG determination protocol. After incubation with the NaSCN concentrations, spectrophotometric readings at 652 nm were converted into immunoglobulin-binding percentage with respect to the values obtained at 0 M NaSCN. Avidity index (AI) is defined as the NaSCN concentration that reduce 50% of immunoglobulin-binding.

#### HLA genotyping (Table S2)

DNA extraction from blood spots of people from endemic malaria areas was carried out following the protocol InstaGene Whole Blood kit (Bio-Rad Laboratories) (31). Briefly, blood spots on Whatman protein saver cards were soaked overnight in a tube containing 100 µL of phosphate-buffered saline (PBS) at 4°C. The tube was centrifuged at 11,000 x g for two minutes. After discarding the supernatant, 100 µL of PBS was added to wash the sediment similarly. The sediment was vortexed for 15 seconds and incubated with 60 µL of InstaGene Matrix CHELEX (Bio-Rad) for 4 minutes at 100ºC. After vortexing again, the tube was centrifuged at 11,000 x g for one minute and the supernatant was aspirated to be purified once again as described above with InstaGene Matrix. The supernatant was then used for the PCR. Aliquots of these DNA samples (20 µl) were taken and HLA-typed by the LABType SSO Class II DRB1 (One Lambda) kit.

For peptide selection, we sought to identify 15-mer peptides in the selected five antigens that could potentially be detected by B-cells and presented by HLA II molecules incorporating HLA-DRB1 alleles. Peptide presentation by HLA-II molecules was assessed after peptide-binding predictions to a panel of HLA-DRB1 molecules expressed by the malaria patient cohort using NetMHCIIpan (2). Only high binders were considered. Subsequently, we selected the minimum set of peptides that for exhibiting promiscuous HLA II peptide-binding covered the all the HLA-DRB1 alleles in the cohort.

**Fig. S1.**

A

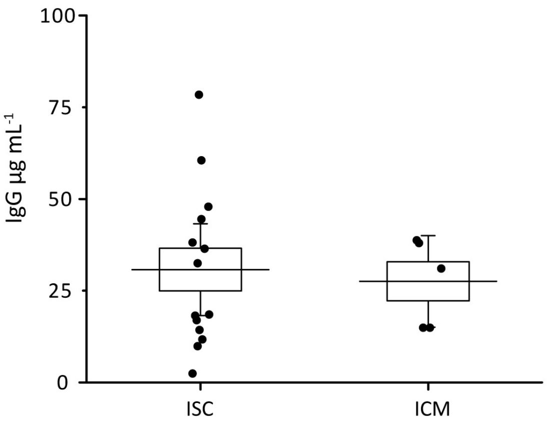


B

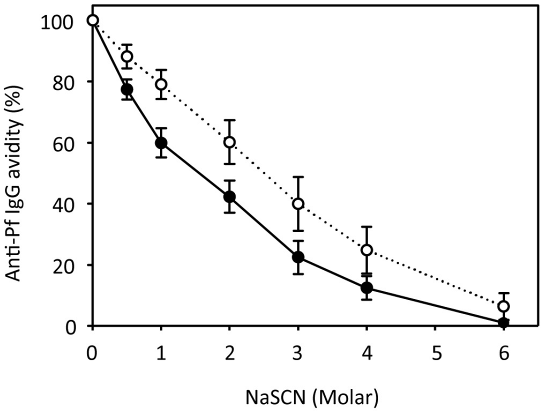


**Figure S1. Concentration of anti-P. falciparum IgG antibody and IgG avidity in sera of imported malaria patients.** A) IgG concentrations obtained by ELISA using immobilized *P. falciparum* total protein as target. Each dot is the IgG concentration of a patient. Horizontal line shows the mean value covered with a box depicting the mean standard error. Whiskers shows the 95% confidence interval. B) IgG avidity. Plot shows mean binding values of sera IgG to total *P. falciparum* protein at different NaSCN concentrations. Open circles are mean values of ISC group; filled circles are mean values of ICM group; bars: standard error. P-value differences = 0.027 at 2 M NaSCN.

**Figure S2**

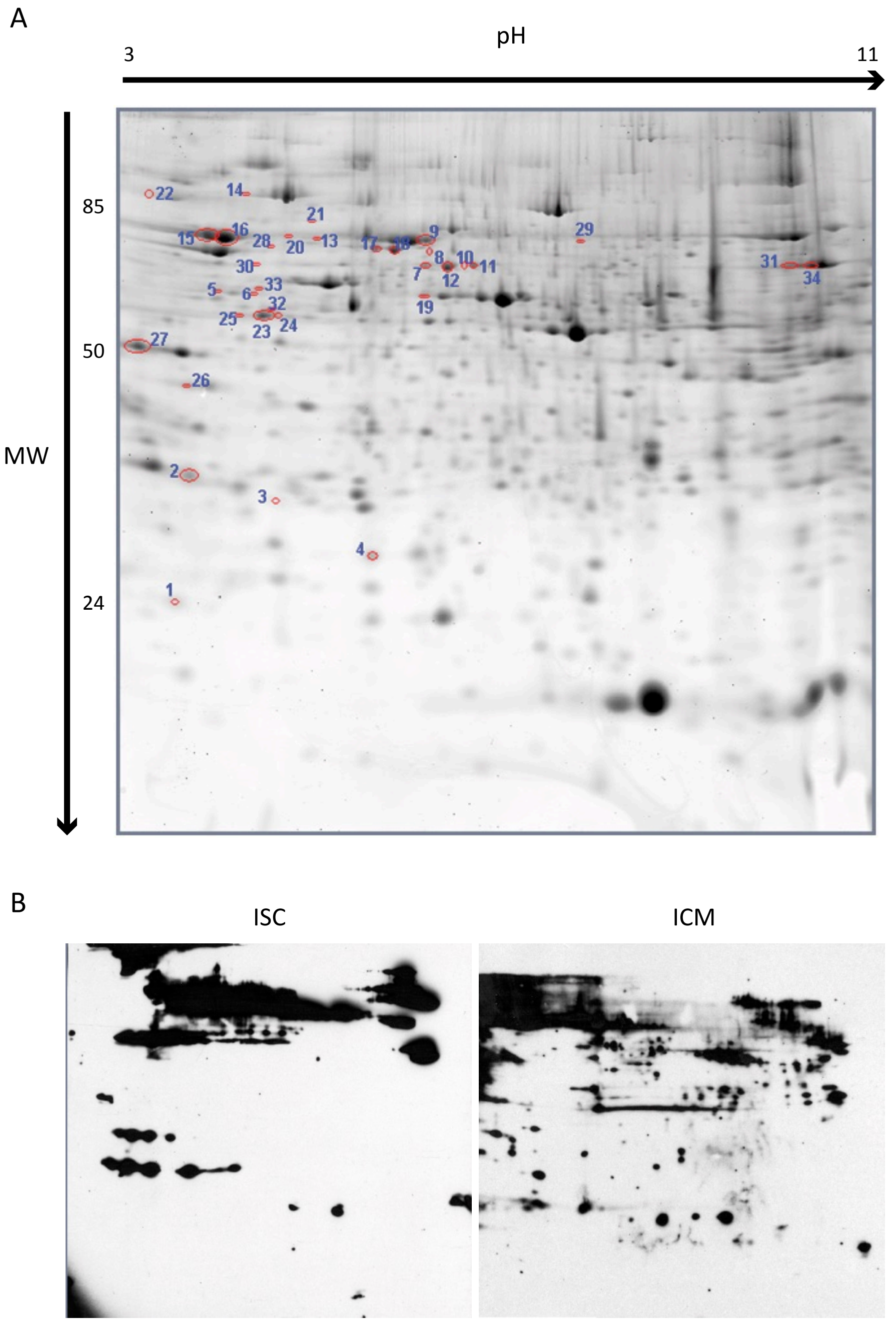


**Figure S2. Two-dimensional electrophoresis profile of *P. falciparum* crude extracts stained with SyBR Pro Ruby and the common IgG reactive proteins detected with sera from imported malaria patients.** A) Total *P. falciparum* proteins separated by 2D gel electrophoresis using a linear pI 3–11 1st dimension. Gels were SYPRO Ruby stained. B) Representative replica gels western blotted with patient sera from ISC and ICM groups. Spots that were recognized by at least 30% of patient sera on parallel western blots were selected for identification (circled in red at panel A). Spots whose identity was determined are numbered in panel A and depict the proteins described in Table 1. See Material and Methods section in main text.

**Table S1.** Demographics, parasitemia and IgG values of imported malaria sera samples used in this study.

| Group[a] | Sample Id. | Sex | Age (years) | *Place of origin* | *Pf-specific IgG µg/mL* |
|---|---|---|---|---|---|
| ISC | 1517 | M | 59 | Equatorial Guinea | 61 |
| | 5517 | M | 28 | Unknown | 12 |
| | 38708 | F | 36 | Equatorial Guinea | 17 |
| | 42350 | F | 45 | Equatorial Guinea | 10 |
| | 44270 | M | 53 | Equatorial Guinea | 18 |
| | 46130 | F | 64 | Equatorial Guinea | 79 |
| | 48751 | F | 57 | Equatorial Guinea | 38 |
| | 48790 | F | 45 | Equatorial Guinea | 32 |
| | 5637 | M | 31 | Cameroon | 2 |
| | 39732 | F | 40 | Equatorial Guinea | 19 |
| | 42098 | M | 73 | Equatorial Guinea | 14 |
| | 49039 | M | 45 | Equatorial Guinea | 36 |
| | 52746 | M | 38 | Equatorial Guinea | 45 |
| | 52899 | F | 69 | Equatorial Guinea | 48 |
| ICM | 6031 | M | 34 | Conakry Guinea | 15 |
| | 35421 | F | 29 | Equatorial Guinea | 39 |
| | 4827 | M | 23 | Cameroon | 31 |
| | 5256 | F | 50 | Undeclared | 15 |
| | 6032 | M | 18 | Senegal | 38 |

[a] ISC: Imported subclinical malaria group, parasitemia <0.1%; ICM: Imported clinical malaria group.

**Table S2.** Demographics, parasitemia and IgG values of sera samples collected at Breman-Asikuma from adults used in this study.

| Group[a] | Sample Id. | Sex | Age (years) | *Pf*-specific IgG µg/mL | Anti-*Pf* IgG High/Low[b] |
|---|---|---|---|---|---|
| ASC | 284 | F | 28 | 4 | L |
| | 64* | F | 25 | 8 | L |
| | 88* | F | 30 | 10 | L |
| | 135 | M | 26 | 10 | L |
| | 253 | M | 26 | 16 | L |
| | 25 | F | 33 | 22 | L |
| | 296* | M | 27 | 22 | L |
| | 86 | F | 31 | 25 | L |
| | 61* | F | 17 | 26 | L |
| | 40 | M | 30 | 45 | L |
| | 225 | F | 26 | 59 | L |
| | 259 | F | 29 | 63 | H |
| | 288* | F | 28 | 63 | H |
| | 145 | M | 35 | 73 | H |
| | 22* | M | 21 | 77 | H |
| | 66 | M | 30 | 84 | H |
| | 60 | M | 28 | 85 | H |
| | 62 | F | 30 | 102 | H |
| | 300* | M | 29 | 160 | H |
| | 307 | F | 33 | 225 | H |
| | 193* | F | 25 | 692 | H |
| ANP | 281* | F | 26 | 14 | L |
| | 310 | M | 40 | 14 | L |
| | 230 | M | 23 | 17 | L |
| | 290* | M | 20 | 20 | L |
| | 100 | M | 28 | 23 | L |
| | 314 | M | 28 | 27 | L |
| | 125* | F | 19 | 39 | L |
| | 141* | M | 22 | 52 | L |
| | 9* | F | 25 | 65 | H |
| | 247 | M | 21 | 67 | H |
| | 132 | F | 26 | 72 | H |
| | 302 | F | 22 | 84 | H |
| | 144 | F | 26 | 89 | H |
| | 209* | F | 20 | 105 | H |
| | 270* | M | 19 | 117 | H |
| | 271 | M | 32 | 130 | H |
| | 4* | F | 26 | 145 | H |

[a] ASC: Adult subclinical malaria group, parasitemia <0.1%; ANP: undetected parasitemia group
[b] Cut off for Hig/Low IgG content 60 µg $mL^{-1}$
* The samples labeled with asterisk were selected for the validation of the 20-mer immunoreactive epitopes (Figure 3).

**Table S3.** Demographics, parasitemia and IgG values of sera samples collected at Breman-Asikuma from children under 5 y.o. used in this study.

| Group[a] | Sample Id. | Age (months) |
|---|---|---|
| CCM | 91 | 60 |
| | 210 | 18 |
| | 71 | 36 |
| | 297 | 18 |
| | 77 | 12 |
| | 95 | 48 |
| | 124 | 36 |
| CSC | 260 | 13 |
| | 298 | 18 |
| | 10 | 60 |
| | 94 | 60 |
| | 303 | 48 |
| | 215 | 24 |
| | 167 | 36 |
| CNP | 201 | 48 |
| | 208 | 1.5 |
| | 150 | 7 |
| | 105 | 8 |
| | 265 | 36 |
| | 53 | 48 |

[a] CCM: Children Clinical Malaria group, parasitemia >0.1%; CSC: Children Subclinical Malaria group, parasitemia <0.1%; CNP: Children with undetected parasitemia

**Table S4.** HLA genotypes of sera samples collected at Breman-Asikuma from adults used in this study.

| Group[a] | Sample Id. | Assigned Allele Pair | Assigned Sero |
|---|---|---|---|
| ASC | 284 | DRB1*08:04 DRB1*09:01 | DR8 DR9 |
| | 64 | DRB1*07:01 DRB1*09:02 | DR7 DR- |
| | 88 | U.D. | U.D. |
| | 135 | DRB1*10:01 DRB1*13:02 | DR10 DR13 |
| | 253 | DRB1*08:04 DRB1*14:01 | DR8 DR14 |
| | 25 | U.D. | U.D. |
| | 296 | DRB1*07:01 DRB1*08:04 | DR7 DR8 |
| | 86 | DRB1*13:02 DRB1*13:24 | DR13 DR- |
| | 61 | DRB1*03:02 DRB1*04:05 | DR18 DR4 |
| | 40 | DRB1*01:02 DRB1*14:123 | DR1 DR- |
| | 225 | U.D. | U.D. |
| | 259 | DRB1*03:02 DRB1*13:02 | DR18 DR13 |
| | 288 | DRB1*11:01 DRB1*13:02 | DR11 DR13 |
| | 145 | DRB1*08:01 DRB1*11:04 | DR8 DR11 |
| | 22 | DRB1*07:01 DRB1*14:01 | DR7 DR14 |
| | 66 | U.D. | U.D. |
| | 60 | U.D. | U.D. |
| | 62 | DRB1*10:01 DRB1*13:02 | DR10 DR13 |
| | 300 | U.D. | U.D. |
| | 307 | U.D. | U.D. |
| | 193 | U.D. | U.D. |
| ANP | 281 | DRB1*01:02 DRB1*03:02 | DR1 DR18 |
| | 310 | U.D. | U.D. |
| | 230 | U.D. | U.D. |
| | 290 | DRB1*03:02 DRB1*15:03 | DR18 DR15 |
| | 100 | DRB1*01:02 DRB1*08:04 | DR1 DR8 |
| | 314 | U.D. | U.D. |
| | 125 | DRB1*10:01 DRB1*13:01 | DR10 DR13 |
| | 141 | DRB1*07:01 DRB1*13:03 | DR7 DR13 |
| | 9 | DRB1*01:02 DRB1*13:02 | DR1 DR13 |
| | 247 | DRB1*01:02 DRB1*08:04 | DR1 DR8 |
| | 132 | DRB1*09:01 DRB1*16:21N | DR9 DR"Blank" |
| | 302 | DRB1*11:01 DRB1*15:03 | DR11 DR15 |
| | 144 | DRB1*07:01 DRB1*08:04 | DR7 DR8 |
| | 209 | U.D. | U.D. |
| | 270 | DRB1*15:03 DRB1*16:02 | DR15 DR16 |
| | 271 | DRB1*03:02 DRB1*13:02 | DR18 DR13 |
| | 4 | DRB1*01:02 DRB1*04:01 | DR1 DR4 |

[a] ASC: Adult subclinical malaria group, parasitemia <0.1%; ANP: undetected parasitemia group
U.D.: Undetermined

**REFERENCES**


1. Pullen GR, Fitzgerald MG, & Hosking CS (1986) Antibody Avidity Determination by Elisa Using Thiocyanate Elution. *J Immunol Methods* 86(1):83-87.

2. Nielsen M, Lundegaard C, Blicher T, Peters B, Sette A, Justesen S, Buus S & Lund O (2008) Quantitative predictions of peptide binding to any HLA-DR molecule of known sequence: NetMHCIIpan. *PLoS Comput Biol.* 4(7):e1000107. doi: 10.1371/journal.pcbi.1000107.
